# SETDB1 prevents TET2-dependent activation of IAP retroelements in naïve embryonic stem cells

**DOI:** 10.1101/193854

**Authors:** Özgen Deniz, Lorenzo de la Rica, Kevin C. L. Cheng, Dominik Spensberger, Miguel R. Branco

## Abstract

**Background:** Endogenous retroviruses (ERVs), which are responsible for 10% of spontaneous mouse mutations, are kept under control via several epigenetic mechanisms. The H3K9 histone methyltransferase SETDB1 is essential for ERV repression in embryonic stem cells (ESCs), with DNA methylation also playing an important role. It has been suggested that SETDB1 protects ERVs from TET-dependent DNA demethylation, but the relevance of this mechanism for ERV expression remains unclear. Moreover, previous studies have been performed in primed ESCs, which are not epigenetically or transcriptionally representative of preimplantation embryos.

**Results:** We used naïve ESCs to investigate the role of SETDB1 in ERV regulation and, in particular, its relationship with TET-mediated DNA demethylation. Naïve ESCs show an increased dependency on SETDB1 for ERV silencing when compared to primed ESCs, including at the highly mutagenic intracisternal A particles (IAPs). We found that, in the absence of SETDB1, TET2 activates IAP elements in a catalytic-dependent manner. Surprisingly, however, TET2 does not drive changes in DNA methylation levels at IAPs, suggesting that it regulates these transposons indirectly. Instead, SETDB1 depletion leads to a TET2-dependent loss of H4R3me2s, which is indispensable for IAP silencing during epigenetic reprogramming.

**Conclusions:** Our results demonstrate a novel and unexpected role for SETDB1 in protecting IAPs from TET2-dependent histone arginine demethylation.

## Background

Endogenous retroviruses (ERVs) are retroelements bearing long terminal repeats (LTRs) and constitute approximately 10% of the mouse genome [1]. Whilst most ERVs are inactive, a subset of these genetic parasites retain their transposition ability and therefore pose a threat to genome integrity [2]. Indeed, around 10% of mouse spontaneous mutations arise as a direct result of ERV insertions [3] and insertional mutagenesis by ERVs is frequently associated with murine cancers [4,5]. Therefore, numerous transcriptional and post-transcriptional host mechanisms have evolved to supress ERV activity. DNA methylation (or 5-methylcytosine; 5mC) plays an essential role in ERV repression in postimplantation embryos and male germ cells [6,7]. However, during early preimplantation and primordial germ cell (PGC) development, the genome undergoes genome-wide DNA demethylation [8–12] and additional mechanisms are required to ensure ERV silencing. Indeed, ERV silencing in embryonic stem cells (ESCs) is largely dependent on post-translational modification of histones, in particular methylation at H3K9. Removal of the H3K9me3 histone methyltransferase SETDB1 and its co-repressor, KRAB-associated protein (KAP1, also known as TRIM28), leads to significant activation of ERVs in ESCs [13–15] and PGCs [16]. Interestingly, 5mC and H3K9me3 regulate largely non-overlapping subsets of ERVs in ESCs, with the notable exception of intracisternal A particles (IAPs), whose silencing depends on the synergistic action of both epigenetic marks [14,15].

IAP elements are relatively resistant to DNA demethylation during epigenetic reprogramming [6, 16–18], which is presumably a host defense mechanism against these highly mutagenic ERVs. Maintenance of methylation at IAP and imprinting control regions is driven by the G9a/GLP complex, which recruits *de novo* DNA methyltransferases (DNMTs) independent of its H3K9 methyltransferase activity [19–21]. On the other hand, H3K9me2-enriched regions are refractory to demethylation during epigenetic reprogramming [22] via recruitment of the DNMT1 chaperone NP95/UHRF1 [23,24].

A role for H3K9me3 in protecting ERVs from TET-mediated DNA demethylation has also been proposed in ESCs [25]. TET enzymes oxidize 5mC into 5- hydroxymethylcytosine (5hmC) and other oxidative derivatives as part of an active DNA demethylation pathway [26–28]. We have previously shown that TET1 binds to multiple retroelements in ESCs, and that both TET1 and TET2 help to maintain LINE- 1 elements in a hypomethylated state [29]. At LTR elements such as IAPs, however, it has been shown that loss of SETDB1 enables TET1 binding, concomitant with an accumulation of 5hmC at these sites [25]. However, this resulted in only very subtle DNA methylation changes, and it remains unknown whether these alterations affect the expression of IAP elements and other ERVs. Moreover, TET enzymes may act on ERVs via non-catalytic pathways, similar to what we observed in LINE-1 elements [29]. Finally, previous studies were performed using primed ESCs grown under standard serum-containing conditions, which are highly methylated and express high levels of the *de novo* methyltransferases DNMT3A and DNMT3B [17,18]. These conditions may counteract and mask the catalytic activity of TET enzymes at IAPs and other ERVs. Global DNA methylation can be induced *in vitro* by growing ESCs under the so-called 2i conditions, which more closely resemble inner cell mass cells, driving a naïve pluripotent state [17,30].

Here we investigated the role of SETDB1 in the regulation of ERVs in naïve ESCs and its relationship with TET-mediated DNA demethylation. We find that SETDB1 has a markedly more prominent role in ERV silencing in naïve cells compared to primed cells, including at IAP elements. The catalytic activity of TET2 contributes to IAP activation upon SETDB1 depletion, but surprisingly this is not linked to DNA methylation changes at IAPs. We show that instead TET2 drives a loss of the repressive H4R3me2s mark at IAPs.

## Results

### SETDB1 safeguards ERV silencing in naïve ESCs

To investigate the role of SETDB1 in ERV silencing in naïve ESCs, we switched serum-grown (i.e., primed) E14 ESCs to 2i culture conditions. Using deep sequencing of PCR amplicons from oxidative bisulfite (oxBS)-treated DNA [31,32], we first confirmed that 5mC levels were substantially lower in naïve versus primed ESCs at multiple ERVs, including IAP elements (Additional File 1: Fig. S1A). RLTR4/MuLV elements were already hypomethylated in primed ESCs and showed only a small decrease in 5mC levels in naïve ESCs. 5hmC levels were generally low and similar between both culture conditions (Additional File 1: Fig. S1B). In line with previous reports [33], transcript levels of these ERVs were not significantly higher in naïve ESCs compared to primed ESCs, suggesting that other mechanisms compensate the loss of DNA methylation to protect the genome against the activity of ERVs (Additional File 1: Fig. S1C).

To identify SETDB1 targets in naïve ESCs, we depleted SETDB1 by lentiviral delivery of shRNAs (Fig. 1A) and performed RNA-seq on three independent biological replicates. For comparison, we conducted the same experiment in primed ESCs. Using an “inclusive mapping” strategy that harvests information from non-uniquely aligned reads (see Material and methods), we identified classes of repetitive elements that were differentially expressed by more than 2-fold (p<0.05, DESeq2). As expected, SETDB1 depletion in naïve ESCs led to the upregulation of many repeat classes (n=104), the vast majority of which were ERVs (Fig. 1B). Notably, about half of these repeat classes (n=55) were exclusively upregulated in naïve cells and not in primed cells (Additional File 2). These naïve-specific repeats included MERVL, LINE-1 and VL30 elements, amongst several others. In contrast, only 8 repeat classes were significantly upregulated exclusively in primed ESCs (Additional File 2).

**Figure 1.**
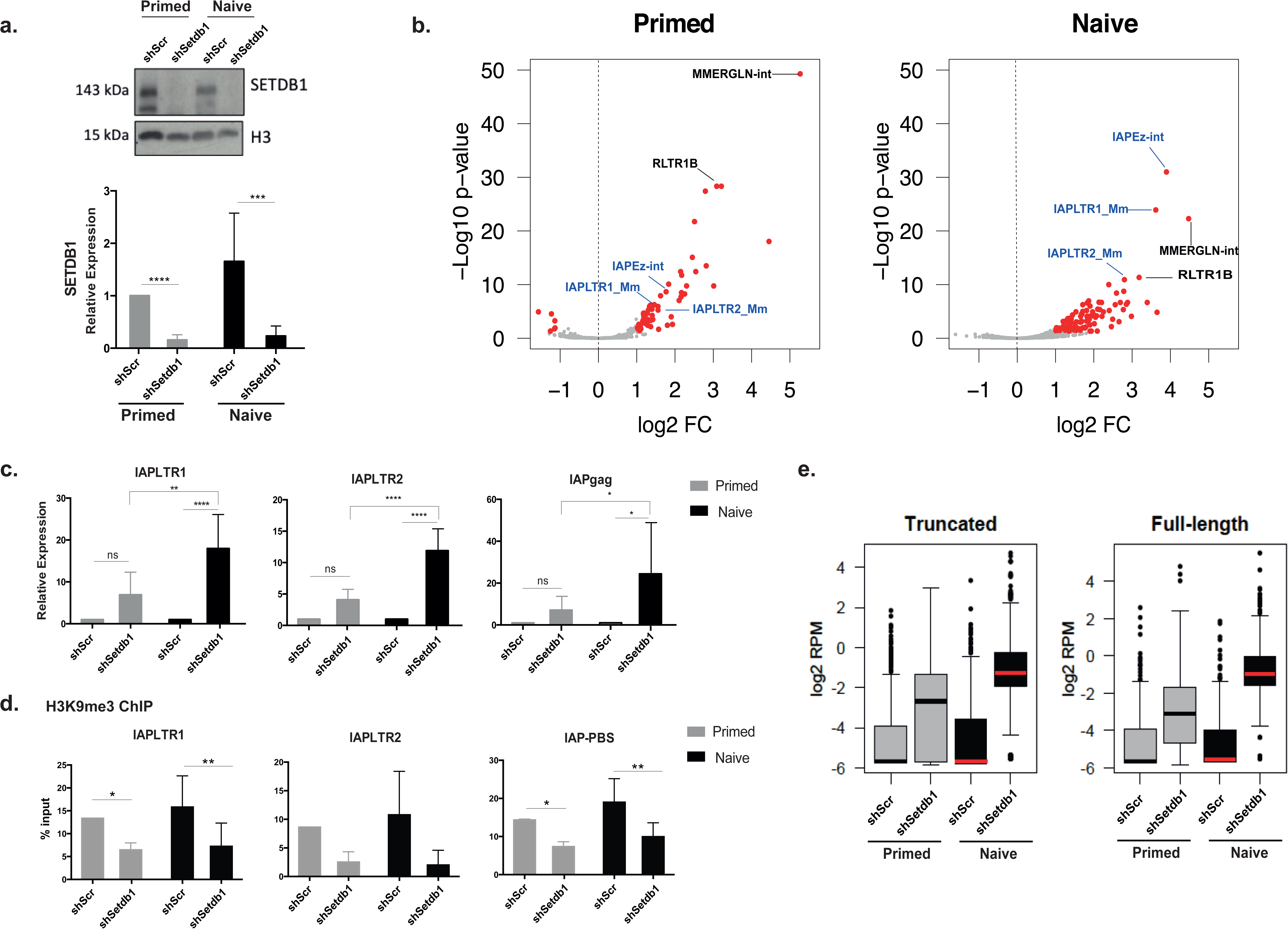
SETDB1 is pivotal for ERV silencing in naïve ESCs. **a.** Western blot (representative data from n=2) and RT-qPCR analyses (n=7) show SETDB1 depletion by lentiviral delivery of shRNAs (ANOVA with Tukey’s multiple comparison test, *** p<0.001, **** p<0.0001). **b.** RNA-seq data using inclusive mapping from primed and naïve cells was overlayed with the RepeatMasker annotation to determine repeat classes that are differentially expressed upon SETDB1 removal (highlighted in red, n=3). **c.** Analysis of IAP expression by RT-qPCR in SETDB1 KD primed and naïve ESCs. The error bars show s.d. (n=6-12; ANOVA with Tukey’s multiple comparison test, * p<0.05, **p<0.01, **** p<0.0001). **d.** H3K9me3 ChIP- qPCR at IAPs upon SETDB1 loss (ANOVA with Tukey’s multiple comparison test, * p<0.05, **p<0.001). Data are shown as mean ± s.d. of seven independent experiments. **e.** Expression data from uniquely mapped RNA-seq reads at truncated and full-length (>5kb) IAP elements.

Intriguingly, although IAP elements were deregulated in both culture conditions, they were substantially more activated in naïve ESCs when compared to primed ESCs (Fig. 1B, Additional File 2). We validated these observations using quantitative reverse transcription polymerase chain reaction (RT-qPCR), which confirmed that IAPEz upregulation was more pronounced in naïve cells (Fig. 1C), even though SETDB1 depletion led to a substantial loss of H3K9me3 at these elements in both conditions (Fig. 1D). Similar results were obtained upon KAP1 depletion (Additional File 1: Fig. S1D), as expected from the dependency of SETDB1 binding on KAP1 [13,15]. We then asked what fraction of IAP copies underwent this skewed IAP activation in SETDB1-depleted naïve cells. We analysed data from uniquely mapped RNA-seq reads and found that, out of 1,009 IAPs with detectable RNA-seq signal, 681 (67%) were found to be >10-fold upregulated upon SETDB1 depletion in naïve ESCs, whereas in primed cells there were only 257 (25%). Notably, this pattern was seen in both full-length and truncated elements (Fig. 1E), showing that SETDB1 removal leads to increased activation at the majority of mappable IAP elements in naïve ESCs compared to primed ESCs.

All together, these data show that SETDB1 plays a more prominent role in ERV silencing in naïve ESCs when compared to primed ESCs, including at IAP elements.

### IAP activation upon SETDB1 depletion depends on TET2 activity

In naïve ESCs the role of SETDB1 in ERVs suppression could be particularly critical for genome integrity due to the hypomethylated state of these transposons. Additionally, SETDB1 could protect ERVs from further DNA demethylation, by preventing binding of TET enzymes [25]. However, it remains unclear to what extent TET activity affects ERV methylation and expression. To address this question, we first performed ChIP-qPCR on WT and TET-depleted ESCs, which revealed that both TET1 and TET2 specifically bind IAP elements at the LTR and primer binding site (PBS; where KAP1 is recruited) regions in both primed and naïve ESCs (Fig. 2A).

**Figure 2.**
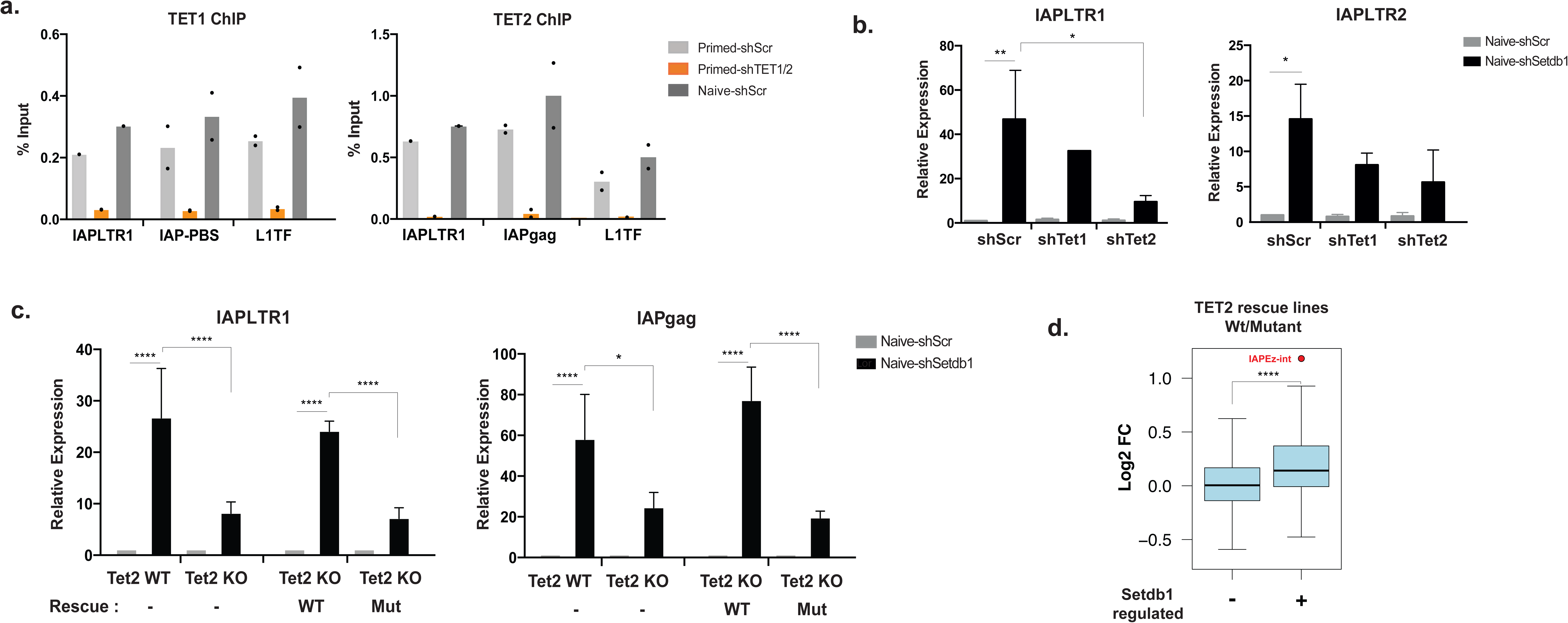
TET2 drives IAP expression in the absence of SETDB1. **a.** ChIP-qPCR data showing TET1 and TET2 enrichment at IAPs and L1 elements in primed (in the presence and absence of TETs) and naïve ESCs. Data points from two independent experiments are shown. **b.** Expression of IAPLTR1 and IAPLTR2 in SETDB1 and TET1 or TET2-depleted cells in naïve ESCs (n=2; one-way ANOVA, Dunnet’s multiple comparison test, * p<0.05, **p < 0.01). Each bar represents the mean ± s.d. **c.** RT-qPCR analysis of *Tet2* WT, KO and rescue cell lines at IAPs in naïve ESCs (n=3-4; ANOVA with Tukey’s multiple comparison test, * p<0.05, **** p<0.0001). The results are presented as mean ± s.d. **d.** Fold-change in expression between TET2 wildtype and mutant rescue lines for different repeats classes, depending on whether they are repressed by SETDB1 or not (n=3, t-test).

Notably, the enrichment of both TET enzymes at IAPs was similar to that seen at LINE-1 elements, which we have previously shown to undergo TET-mediated DNA demethylation [29]. To test whether TET enzymes were involved in the activation of IAPs, we performed *Tet1* or *Tet2* knockdown (KD) in SETDB1-depleted naïve ESCs. Our RT-qPCR analyses revealed that removal of TET2 markedly reduced IAPLTR1 activation (and IAPLTR2 to a lesser extent) in SETDB1-depleted cells, whereas this effect was milder upon *Tet1* KD (Fig 2B). We also generated *Tet2* knockout (KO) ESCs (Additional File 1: Fig. S2A,B) wherein, similarly to *Tet2* KD cells, upregulation of IAPLTR1, as well as the coding *gag* region, was diminished when compared to *Tet2* wild-type (WT) naïve ESCs (Fig. 2C). Experiments involving depletion of both TET1 and TET2 showed that loss of TET2 alone was sufficient to maximally impair IAP activation (Additional File 1: Fig. S2C).

We then asked whether the effect of TET2 is dependent on its 5mC-oxidising catalytic activity. For this purpose, we used *Tet2* KO ESCs to establish stable cells lines expressing either WT TET2 protein or a catalytically inactive mutant version of the enzyme. Western blot analyses revealed that both wild-type and mutant proteins were expressed at similar levels (Additional File 1: Fig. S2D). Upon SETDB1 depletion, naïve ESCs expressing the WT *Tet2* construct displayed similar activation of IAPLTR1 and *gag* region to what was seen in *Tet2* WT cells, effectively rescuing the loss of TET2 (Fig. 2C). In contrast, cells expressing the catalytic mutant TET2 failed to upregulate IAPs any further than what was seen in SETDB1-depleted *Tet2* KO cells (Fig. 2C). These results show that the catalytic activity of TET2 contributes to the activation of IAPs upon SETDB1 depletion. To test whether other ERVs were targeted by the same mechanism, we performed RNA-seq on both TET2 rescue lines in a SETDB1-depleted context. Strikingly, comparison of ERV expression between WT and mutant TET2 rescue lines yielded only IAPEz elements as significantly differentially expressed. However, as a group, SETDB1-regulated repeats displayed higher expression levels in WT versus mutant TET2 rescue lines upon SETDB1 depletion, a tendency that was not seen at repeats that are not targeted by SETDB1 (Fig. 2D).

We also performed experiments in primed ESCs, wherein *Tet2* KO had no effect on IAP upregulation upon SETDB1 depletion (Additional File 1: Fig. S2E). On the other hand, both RT-qPCR and RNA-seq data showed that overexpression of wild-type TET2 could also drive an increase in IAP activation in primed ESCs (Additional File 1: Fig. S2E).

Overall, these results reveal that the activation of IAPs seen upon SETDB1 loss partly depends on the catalytic activity of TET2.

### SETDB1 does not protect IAPs from TET-mediated DNA demethylation

The contribution of TET2 catalytic activity to IAP activation in SETDB1-depleted naïve cells suggests that SETDB1 protects IAPs from oxidation-driven DNA demethylation. To directly address this hypothesis, we measured 5mC and 5hmC levels at IAPs using oxBS. Surprisingly, we did not observe any significant changes in 5mC levels in SETDB1-depleted naïve ESCs both at the LTR and PBS regions (Fig. 3A). IAP 5hmC levels remained low after SETDB depletion and only the PBS region displayed a small increase in 5hmC levels (Supplementary Fig.3A). In line with these observations, we found that TET2 binding to IAPs was not enhanced by the loss of SETDB1 in naïve ESCs (Fig. 3B).

**Figure 3.**
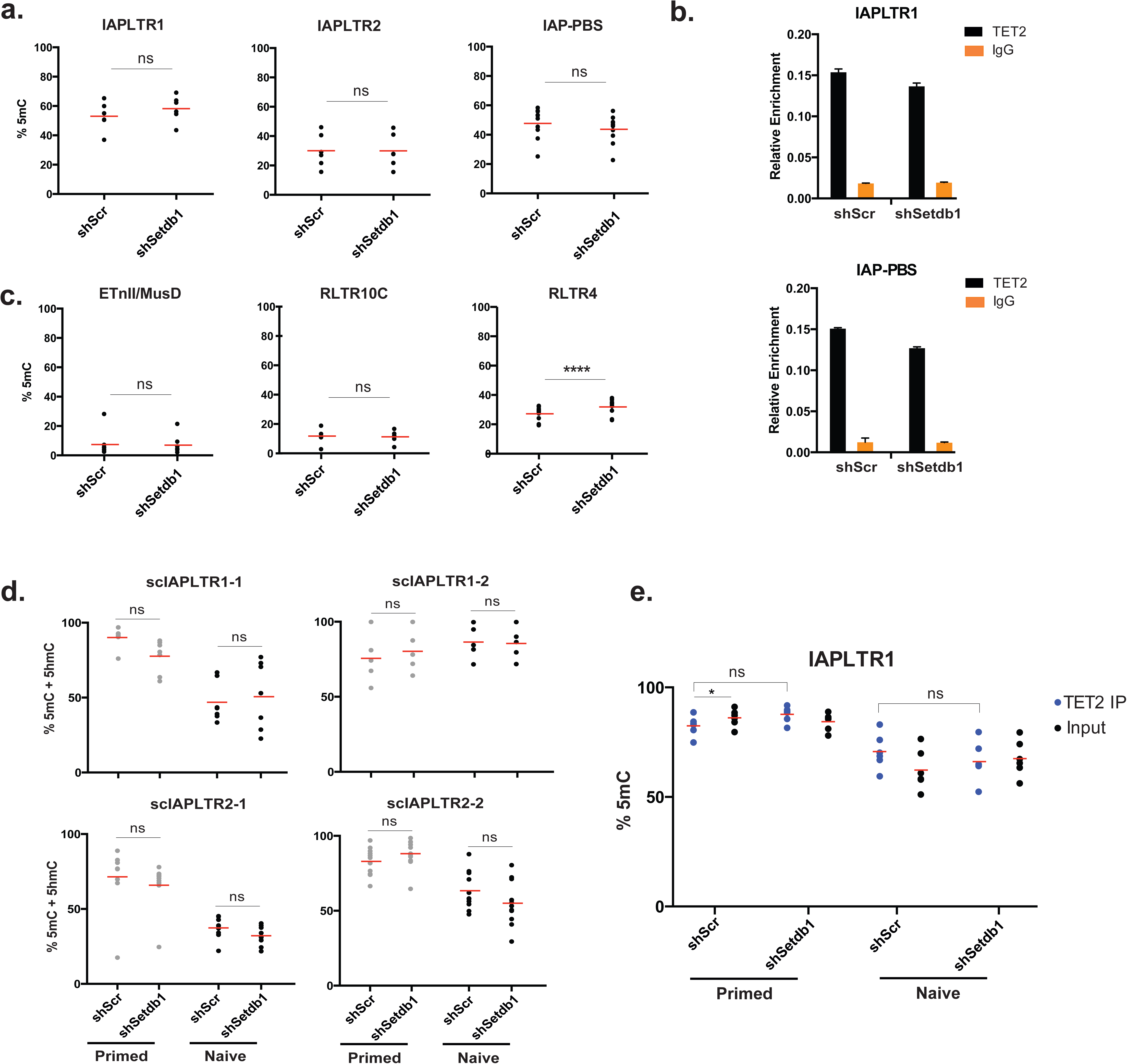
TET2 does not affect DNA methylation levels at IAPs in naïve ESCs. **a.** 5mC levels using oxBS at IAPs upon SETDB1 knockdown in naïve ESCs; each data point represents the average value from two biological replicates at a given CpG within the amplicon. **b.** Normalised enrichment of TET2 using a spike-in control at IAPLTR1 and the PBS region in naïve ESCs (representative replicate from n=3) **c.** 5mC levels at additional ERVs upon SETDB1 knockdown in naïve ESCs (ANOVA with Tukey’s multiple comparison test, **** p<0.0001). **d.** 5mC+5hmC levels using BS at individual IAP copies (scIAPs) upon SETDB1 depletion, in primed and naïve ESCs. **e.** 5mC levels at TET2-bound IAPLTR1 elements; each data point represents the value at a given CpG within the amplicon (ANOVA with Tukey’s multiple comparison test, * p<0.05).

Extending our analyses to other SETDB1-regulated ERVs, we found that ETn/MusD, RLTR10C and VL30 elements also did not display a decrease in 5mC upon SETDB1 depletion in naïve ESCs (Fig. 3C). In contrast, primed ESCs displayed a small but significant loss of 5mC at IAPLTR2, ETnII/MusD and RLTR10C ERVs upon SETDB1 depletion, which is consistent with previous findings [13,25] (Additional File 1: Fig. S3B). However, these small reductions in 5mC levels were not associated with changes in expression, as IAPLTR2 transcripts are not affected in primed *Tet2* KO cells (Additional File 1: Fig. S2E). Furthermore, the lack of 5mC changes in naïve cells suggests that TET2 does not affect ERV expression by driving their demethylation.

We next considered the possibility that 5mC changes may be apparent in copies that are more responsive to SETDB1 depletion, rather than in the pool of IAP copies that are amplified by the consensus primers used above. Based on our RNA-seq data, we designed specific primers for bisulfite sequencing that target three individual IAPLTR1 and two individual IAPLTR2 elements that exhibited high activation upon SETDB1 depletion. Notably, these individual elements show higher levels of TET2 binding compared to a pool of IAP copies, whereas H3K9me3 and TET1 levels are similar (Additional File 1: 3C). Despite displaying higher TET2 levels, the tested IAP copies did not show any significant alterations in DNA methylation in the absence of SETDB1 in either naïve or primed ESCs (Fig. 3D, Additional File 1: Fig. S3D).

To confirm that TET2 did not modulate DNA methylation levels at IAPs, we performed bisulphite sequencing in TET2-depleted naïve cells. As expected, removal of TET2 did not affect methylation levels both for a pool of IAP copies as well as individual copies (Additional File 1: Fig. S3E). Finally, we asked whether the effect of TET2 on DNA methylation is evident only at its target IAP copies. To this end, we analysed methylation levels of DNA that is immunoprecipitated by a TET2 antibody. DNA methylation levels of TET2-bound IAPs were comparable to the whole pool of IAPs (input) in all conditions, and no differences were seen between TET2-bound elements before or after SETDB1 depletion (Fig. 3E).

Taken together, these observations reveal that SETDB1 does not safeguard IAPs from TET-mediated DNA demethylation and that TET2 induces IAP activation by a DNA methylation-independent mechanism.

### TET2 expression is associated with loss of H4R3me2 at IAPs

We hypothesized that the contribution of TET2 to ERV activation in SETDB1- depleted naïve ESCs could be affecting key histone marks, possibly through an indirect mechanism. Therefore, we first asked whether TET2 could contribute to the loss of H3K9me3 seen upon SETDB1 depletion, but found that *Tet2* knockdown did not affect the levels of H3K9me3 in naïve ESCs (Fig. 4A). It was previously shown that TET enzymes can recruit O-GlcNac transferase (OGT) to chromatin in ESCs [34], which in turn targets H3K4 methyltransferase SET1/COMPASS complex [35]. However, our ChIP-qPCR analyses demonstrated that TET2 depletion did not affect H3K4me3 levels at IAPs, which remained very low in all conditions tested (Fig 4A). We also found no differences in the levels of the H3K27me3 repressive mark (Additional File 1: Fig. S4A). IAP elements were also shown to be highly enriched for symmetrical arginine dimethylation at H4R3 (i.e., H4R3me2s) [36,37]. Importantly, removal of the arginine methyltransferase PRMT5 leads to derepression of IAPs in PGCs and blastocysts [37], suggesting that H4R3me2s is a key repressive mark of IAPs during epigenetic reprogramming. Therefore, we next carried out ChIP-qPCR for H4R3me2s and found reduced levels of this mark in SETDB1-depleted ESCs (Fig. 4B). Strikingly, this loss at IAPs was driven by the action of TET2, as the levels of H4R3me2s remained stable upon SETDB1 removal in *Tet2* knockdown cells (Fig. 4B). These results suggest that TET2 contributes to IAP activation in SETDB1-depleted ESCs by modulating the levels of the repressive H4R3me2s mark.

**Figure 4.**
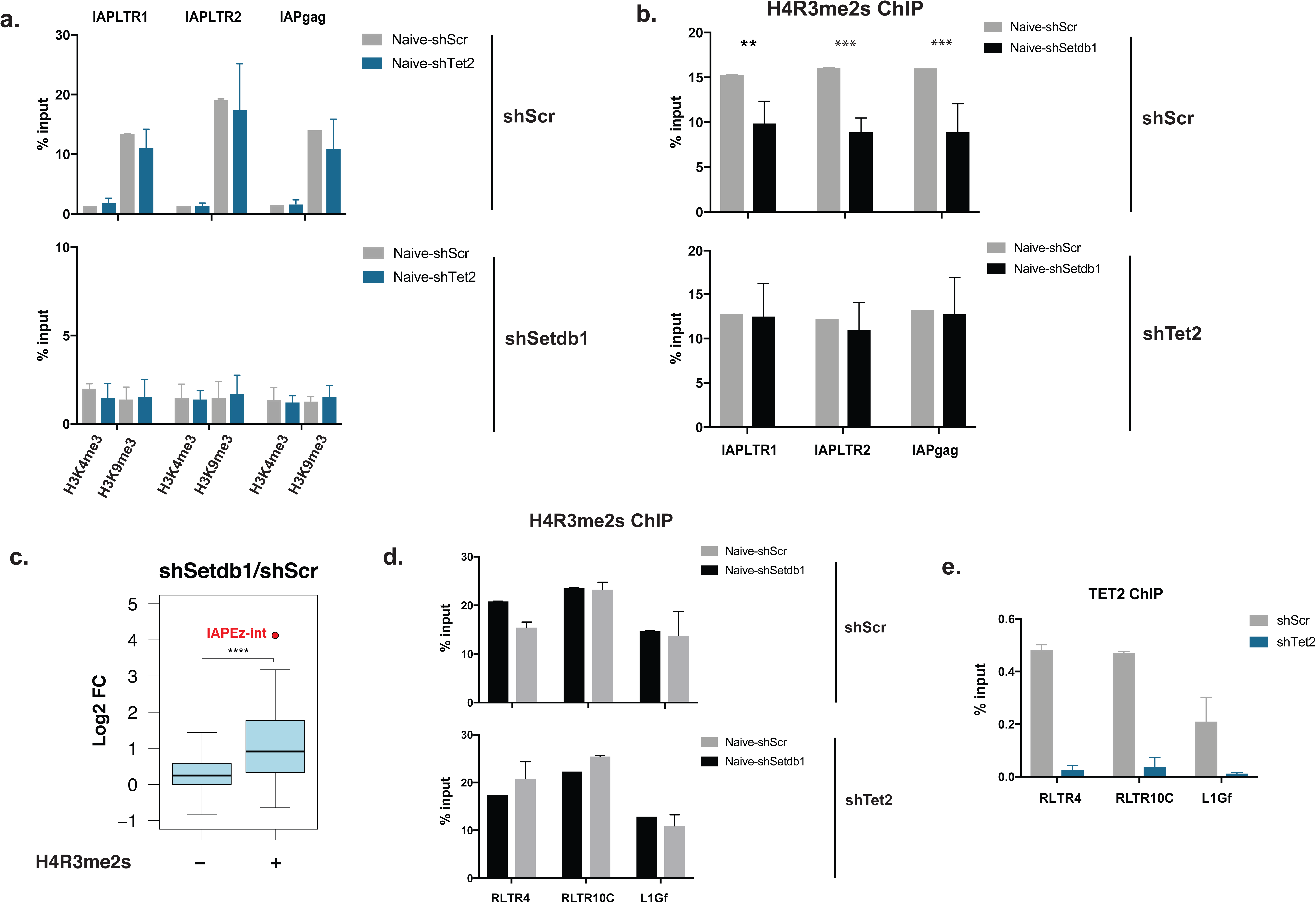
SETDB1 protects IAPs from TET2-dependent loss of H4R3me2s. **a,b.** ChIP-qPCR data for H3K4me3 and H3K9me3 (a) or H4R3me2s (b) at IAPs upon SETDB1 loss in the presence and absence of TET2 (n=2-3; ANOVA with Sidak’s multiple comparison test, **p<0.01, *** p<0.001). **c.** Fold-change in expression upon SETDB1 knockdown for different repeats classes, depending on whether they are enriched for H4R3me2s peaks or not (n=3, t-test). **d,** ChIP-qPCR data for H4R3me2s at additional ERVs upon SETDB1 loss in the presence and absence of TET2 (n=2, each bar represents the mean ± sd). **e.** TET2 enrichment levels at ERVs. Data are shown as mean ± s.d. of two independent experiments.

To test whether a similar mechanism could be responsible for the activation of other ERVs, we mined publicly available ChIP-seq data for H4R3me2s in naïve ESCs [36]. Using uniquely mapped reads, we identified repeat classes that are enriched for H4R3me2s peaks over a random control (Additional File 1: Fig. S4B). Interestingly, H4R3me2s-enriched repeats were preferentially activated upon SETDB1 depletion when compared to non-enriched repeats (Fig. 4C). Using ChIP-qPCR we validated the enrichment of H4R3me2s on three selected SETDB1-regulated transposons (RLTR4/MuLV, RLTR10C and L1Gf) and tested whether, similar to IAPs, SETDB1 depletion led to H4R3me2s loss at these sites. However, none of the tested elements displayed a significant reduction in H4R3me2s levels upon SETDB1 removal (Fig. 4D). This is in line with the fact that, unlike IAPs, the expression of these transposons was not mediated by the catalytic activity of TET2 (Additional File 1: Fig. S4C). These data suggest that TET2-mediated loss of H4R3me2s is specific to IAPs and drives their activation.

Notably, all of the transposons analysed above displayed similar levels of TET2 enrichment (Fig. 4E), showing that TET2 binding is not sufficient to impart a reduction in H4R3me2s levels. This suggest that TET2 activates IAPs in an indirect manner, possibly by regulating the expression of one or more chromatin modifiers that act on a subset of SETDB1-regulated transposons.

## Discussion

Here we have used naïve mouse ESCs to show that SETDB1 protects IAPs from TET2-dependent activation, but that instead of DNA demethylation this involves modulation of H4R3me2s levels at these elements.

We first established that SETDB1 has a more prominent role in ERV silencing in naïve ESCs than in primed ESCs. In contrast, in mouse embryonic fibroblasts ERV supression is largely independent of SETDB1 [13], suggesting that cell differentiation is generally associated with a switch from an H3K9me3-dependent silencing mechanism to a 5mC-dependent one. Our results suggest that such a reciprocal relationship extends further back into naïve pluripotency, where there is a more pronounced requirement for SETDB1-mediated deposition of H3K9me3 for maintaining ERV silencing.

We show for the first time that the catalytic activity of TET2 contributes to IAP activation in SETDB1-depleted naïve ESCs. Unexpectedly, TET2 does not drive DNA demethylation at IAPs, including at individual IAP copies and at TET2-bound IAPs in naïve ESCs (Fig. 4). In contrast, a previous report suggested that in primed ESCs H3K9me3 deposition by SETDB1 protects IAPs from TET-mediated DNA demethylation [25]. Our data in primed ESCs partly supports this (Additional File 1: Fig. S5A), indicating that a potential direct relationship between SETDB1 and TET-mediated DNA demethylation is exclusive to the primed state and seemingly lost in naïve ESCs. Notably, even in primed ESCs, the extent of DNA demethylation is relatively small and is not associated with expression changes. We find that, rather than affecting IAP expression through changes in DNA methylation, TET2 drives a decrease in H4R3me2s levels at IAPs in naïve ESCs (Fig. 4B). It has been shown that in both PGCs and preimplantation embryos, deletion of *Prmt5* leads to loss of H4R3me2s at IAPs, concomitant with their transcriptional activation [37]. Importantly, *Prmt5*-null PGCs display no differences in IAP DNA methylation levels compared to wild-type tissues [37]. Our data in naïve ESCs adds further support to a repressive role of the H4R3me2s mark at IAPs, as other key histone modifications were excluded as the mediators of TET2-dependent IAP activation (Fig. 4A; Additional File 1: Fig. S4A).

Whilst the catalytic activity of TET2 does not affect DNA methylation directly at IAPs, it remains formally possible that TET2 oxidises methylated proteins or RNA associated with IAP chromatin. However, our data suggest that the action of TET2 on IAPs is likely to be indirect, involving the regulation of genes that in turn control IAP expression. Our RNA-seq data show that neither the arginine methyltransferases *Prmt5* and *Prmt7* nor the putative H4R3me2s demethylase *Jmjd6* are controlled by TET2 activity (Additional File 1: Fig. S4D). It remains to be tested whether other enzymes act *in vivo* to modify H4R3me2s, which could then mediate the activating effect of TET2 on IAPs. Other more indirect scenarios are also possible, such as a TET2-regulated gene that prevents recruitment of the enzymes involved in arginine methylation.

If TET2-mediated demethylation indirectly affects IAP expression, then the same could be true for other hypomethylation models, such as *Dnmt* KO ESCs. This raises questions about reported roles of DNA methylation on IAP expression, namely its synergistic action with H3K9me3 [14,38]. Similarly, *Tet1/Tet3* double KO preimplantation embryos display a downregulation of IAPs [39], but direct evidence for a causal relationship is missing. These considerations highlight the need for future work to harvest the power of epigenetic editing tools to test for direct causal links between ERV methylation and their activation.

## Conclusions

We have demonstrated that SETDB1 has more prominent role in ERV silencing in naïve ESCs than primed ESCs, with the removal of SETDB1 leading to increased upregulation of ERVs. Our data show that activation of IAPs in SETDB1 depleted naïve cells depends on the catalytic activity of TET2. However, surprisingly, TET2 does not play a role in DNA demethylation at IAPs, instead TET2-dependent activation of IAPs is associated with the loss of H4R3me2s repressive mark upon SETDB1 removal. In conclusion, our findings demonstrate a novel role of SETDB1 in protecting IAPs from TET2-dependent loss of H4R3me2s in naïve ESCs.

### Methods

#### Cell lines

E14 ESCs were used for all experiments unless otherwise stated. *Tet1* KO ESCs were a kind gift from Guo-Liang Xu [40]. *Tet2* KO ESCs were generated by targeting exon 14 of *Tet2* with a floxed neomycin resistance cassette (Additional File 1: Fig. S4A). To rescue the expression of TET2, stable cell lines were derived from *Tet2* KO ESCs using a PiggyBac transposon system. The *Tet2* catalytic mutant (H1304Y, D1306A) construct was made from a wild-type clone (a kind gift from Kristian Helin) by site-directed mutagenesis.

### Cell culture and gene knockdown

ESCs were grown in feeder-free conditions using either DMEM-based medium with 15% FBS and 1,000 U/ml ESGRO LIF (Millipore) or in 2i culturing conditions using DMEM F-12 (Gibco 21331) and neurobasal media (Gibco 21102) supplemented with N2 (Life tech. 17502048), B27 (Gibco 17504-044), 1,000 U/ml ESGRO LIF (Millipore), Mek inhibitor (PD0325901) and GSK3b inhibitor (CHIR99021). For shRNA-mediated gene knockdown, ESCs were infected with viral particles carrying pLKO.1 constructs harbouring gene-specific shRNAs from The RNAi Consortium, (shSETDB1: CCCGAGGCTTTGCTCTTAAAT, TRCN0000092975; shTET1: TTTCAACTCCGACGTAAATAT, TRCN0000341848; shTET2: TTCGGAGGAGAAGGGTCATAA, TRCN0000250894) or a non-targeting sequence (shScr: CCTAAGGTTAAGTCGCCCTCGCTC;). After 48 hours, cells were selected with 1μg/ml puromycin or 50μg/ml hygromycin for 3 days. For siRNA-mediated gene knockdown, cells were transfected twice (at day one and three) with Kdm5d-specific siRNAs (Dharmacon siGENOME siRNA Cat. D-054675-02-0002 and Thermofisher Silencer Select siRNA Cat. s74014) or non-targeting siRNA (Dharmacon siGENOME Non-Targeting siRNA #2 Cat.D-001210-02-20) at 125nM using Lipofectamin 3000 (Thermo Scientific, Cat. L3000008). The cells were collected four days after the first transfection and the knockdown was confirmed by RT-qPCR.

### RNA Isolation and RT-qPCR

RNA was extracted using AllPrep DNA/RNA mini kit (Qiagen 80204) and DNAse treated with the TURBO DNA-free™ Kit (Ambion, AM1907). RNA (1 μg) was retrotranscribed using Revertaid Reverse Transcriptase (Thermo Scientific EP0441) and the cDNA was diluted 1/50 for qPCRs using MESA BLUE MasterMix (Eurogenentec, 10-SY2X-03+NRWOUB) on a LightCycler^®^ 480 Instrument II (Roche). A list of primers used can be found in Additional File 3.

### RNA-seq Library Preparation

For analysing the effects of SETDB1 depletion, ribosomal RNA-depleted RNA-seq libraries were prepared from 200-500 ng of total RNA using the low input ScriptSeq Complete Gold Kit (Epicentre). For the TET2 rescue samples, mRNA libraries were prepared using the Dynabeads mRNA purification kit (Life Technologies) and the NEBnext Ultra RNA library prep kit (NEB). Libraries were sequenced on an Illumina NextSeq 500 with single-end 75 bp reads.

### Chromatin Immunoprecipitation

ChIP was performed as described in Latos et al [41], with modifications. For the detection of TET1, TET2 and ATRX, cells were fixed with an initial cross-linking step of 45 minutes with 2 mM Di(N-succinimidyl) glutarate (Sigma-Aldrich Cat. 80424) in PBS at room temperature, followed by a PBS wash and a second fixation step of 12 minutes with 1% formaldehyde in PBS. For histone ChIPs (H4K20me3, H3K9me3, H3, H3K4me3, H4R3me2s) the cells were fixed with % formaldehyde for 12 minutes in PBS. After quenching with glycine, washes and lysis, chromatin was sonicated using a Bioruptor Pico from Diagenode, to an average size of 200-700 bp. Immunoprecipitation was performed using 100 μg of chromatin and 7.5 μg of antibody (TET1, TET2, ATRX) or 20 μg of chromatin and 2.5 μg of antibody (histones). Final DNA purification was performed using the GeneJET PCR Purification Kit (Thermo Scientific. Cat. K0701) and elution in 80 μL of elution buffer. This was diluted 1/10 and analysed by qPCR, using the KAPA SYBR^®^ FAST Roche LightCycler® 480 2X qPCR Master Mix (Kapa Biosistems, Cat. KK4611). A list of primers and antibodies used can be found in Additional Files 3 and 4, respectively.

### Oxidative bisulphite sequencing

Deep sequencing of PCR products from BS- and oxBS-converted DNA was performed as previously described [31]. Briefly, precipitated DNA (without glycogen) was resuspended in water and further purified using Micro Bio-Spin columns (Bio-Rad), after which half of the DNA was oxidised with 15 mM KRuO_4_ (Alpha Aesar) in 0.5 M NaOH for 1 hour. Following bisulphite conversion of both DNA fractions with the EpiTect Bisulfite kit (QIAGEN), a two-step PCR amplification was used: a first PCR amplifies the region of interest and adds part of the sequencing adaptors; a second PCR on pooled amplicons then completes the adaptors and adds sample barcodes, allowing for multiplexing (see primers in Additional File 3). Paired-end sequencing of pooled samples was done using an Illumina MiSeq.

### High-throughput sequencing data processing

Reads were trimmed using Trim_galore! v0.3.3 with default parameters. External ChIP-seq data for H4R3me2s (GEO accession GSE37604) [36] were aligned to the mm9 genome assembly using Bowtie2 v2.1.0 [42] and uniquely aligned reads were extracted for peak detection using MACS2. To identify repeats enriched for H4R3me2s, the number of ChIP-seq peaks overlapping each repeat class were compared with a random control where peaks were shuffled (using bedtools) over mappable regions of the genome. RNA-seq data were aligned to mm9 using Tophat v2.0.9 [43] with -g 1 option, which assigns reads with multiple hits of equal quality to one of those locations at random (i.e., “inclusive mapping”). Raw read counts for each gene or Repeatmasker class were used in DESeq2 for differential expression analysis and for generating normalised gene and repeat expression values. For expression analysis of individual IAP copies, only uniquely mapped reads were used, together with a custom annotation of IAPs which merged same-strand IAP fragments within 100bp into a single element; elements longer than 5 kb were classified as full-length. Only elements with >0.25 RPM in any of the analysed samples were used. OxBS data were aligned with Bismark [44] to a custom genome containing the amplicon sequences; only CpGs covered by at least 100 reads were used to calculate 5mC/5hmC levels.

## Declarations

### Availability of data and material

The datasets generated during the current study (RNA-seq) are available in the GEO repository under the accession number GSE100864. ChIP-seq data for H4R3me2s (GEO accession GSE37604) [36] were downloaded from the NCBI Gene Expression Omnibus.

## Acknowledgements

We thank the Barts Genome Centre for high-throughput sequencing, Guo-Liang Xu for the *Tet1* KO ESCs, Tony Green for the *Tet2* KO ESCs, Jesper Christensen and Kristian Helin for the anti-TET2 antibody and the *Tet2* entry clone, and Paul Hurd and Vardhman Rakyan for critical reading of the manuscript.

## Author’s Contributions

Ö.D. and M.B. designed the study and experiments. Ö.D. performed cell culture, shRNA knockdowns, RT-qPCR, Western blotting, oxBS and ChIP experiments. L.R. performed ChIP experiments. K.C. performed qPCR analyses. D.S. generated the *Tet2* KO ESCs. M.B. performed oxBS, RNA-seq and bioinformatic analyses. Ö.D. and M.B. wrote the manuscript with all other authors.

## Ethics

Not applicable.

## Corresponding author

Correspondence to Miguel R. Branco

